# The FinnBrain Multimodal Neonatal Template and Atlas Collection: T1, T2, and DTI brain templates, and accompanying cortical and subcortical atlases

**DOI:** 10.1101/2024.01.18.576325

**Authors:** Jetro J. Tuulari, Aylin Rosberg, Elmo P. Pulli, Niloofar Hashempour, Elena Ukharova, Kristian Lidauer, Ashmeet Jolly, Silja Luotonen, Hilyatushalihah K. Audah, Elena Vartiainen, Wajiha Bano, Ilkka Suuronen, Isabella L.C. Mariani Wigley, Vladimir S. Fonov, D. Louis Collins, Harri Merisaari, Linnea Karlsson, Hasse Karlsson, John D. Lewis

**Author notes:** **Corresponding author**, Jetro J. Tuulari, Department of Clinical Medicine, FinnBrain Birth Cohort Study University of Turku, Finland.

## Abstract

The accurate processing of neonatal and infant brain MRI data is crucially important for developmental neuroscience, but presents challenges that child and adult data do not. Tissue segmentation and image coregistration accuracy can be improved by optimizing template images and / or related segmentation procedures. Here, we describe the construction of the FinnBrain Neonate (FBN-125) template; a multi-contrast template with T1- and T2-weighted as well as diffusion tensor imaging derived fractional anisotropy and mean diffusivity images. The template is symmetric and aligned to the Talairach-like MNI 152 template and has high spatial resolution (0.5 mm^3^). In addition, we provide atlas labels, constructed from manual segmentations, for cortical grey matter, white matter, cerebrospinal fluid, brainstem, and cerebellum as well as the bilateral hippocampi, amygdalae, caudate nuclei, putamina, globi pallidi, and thalami. We provide this multi-contrast template along with the labelled atlases for the use of the neuroscience community in the hope that it will prove useful in advancing developmental neuroscience, for example, by helping to achieve reliable means for spatial normalization and measures of neonate brain structure via automated computational methods. Additionally, we provide standard co-registration files that will enable investigators to reliably transform their statistical maps to the adult MNI space, which has the potential to improve the consistency and comparability of neonatal studies or the use of adult MNI space atlases in neonatal neuroimaging.

## Introduction

Neonatal and infant brain segmentation remains one of the biggest challenges for neuroscientists. Although multiple segmentation procedures have been developed, used, validated and published as openly available software (Devi et al., 2015; Li et al., 2019; Makropoulos et al., 2018), it may be challenging to map and choose the best available tools that are likely to work across data sets. This is in stark contrast to operating with adult MRI data where already validated software is available. Neonatal MRI images have inconsistent tissue contrast that stem from the initial near absence of myelin-related contrast and its uneven pattern of development during the first year of life. In neonates, in areas with little to no myelin, the white matter is darker than the grey matter on t1-weighted images, and lighter than the grey matter on t2-weighted images. Visually, the neonatal brain has roughly the reverse of the adult contrast in structural MR images. But, in areas showing early myelination, the two tissue classes can be almost indistinguishable. Important advancements in the field have been made by introducing high quality templates and accurate anatomical labels to guide final segmentations that can then aid current and future segmentation algorithms (Oishi et al., 2019; Zöllei et al., 2020).

Currently available neonatal and infant atlases are comprehensively introduced in recent review articles (Dufford et al., 2022; Li et al., 2019; Oishi et al., 2019). One of the reviews also aptly suggests that there may not be a one-size-fits-all atlas for neonates. Crucially, the neonatal period and early infancy are dynamic phases of brain development, and investigators likely benefit from having multiple available atlases (Oishi et al., 2019; Zöllei et al., 2020) and ultimately robust procedures across different stages of brain development (i.e., 4D templates and atlases across several ages). In addition to contributing to the available neonatal atlases, our work is especially motivated by a recent review pointing out that there is a lack of standard template spaces in neonatal / infant neuroimaging studies and that correspondence to adult MNI space would be helpful in supporting comparisons to adult studies, performing meta-analyses, and assuring reproducibility (Dufford et al., 2022). Finally, our review of available neonate atlases indicates that there is paucity of multimodal templates for neonates (Table 1, included as supplement 1 here).

This article describes a new multi-contrast template for the neonate brain, comprised of T1- and T2-weighted data, as well as diffusion tensor imaging (DTI) data in terms of fractional anisotropy (FA) and mean diffusivity (MD) data. We also created accompanying neonatal brain atlases with the majority vote technique using manually defined labels of 21 subtemplates (Acosta et al., 2020). The atlases were created with: 1) gross tissue labels for grey matter, white matter and cerebrospinal fluid (CSF); 2) symmetric labels for grey matter, white matter, cerebrospinal fluid (CSF), brainstem and cerebellum, as well as labels of the bilateral amygdala, hippocampus, caudate, putamen, globus pallidus and thalamus; and 3) corresponding asymmetric labels for left and right hemisphere with FreeSurfer look up table labels (the templates and labels themselves are symmetric). Finally, we provide standard co-registration files to enable standard transforms to adult MNI coordinates for existing and future studies.

## Methods

The study was conducted according to the Declaration of Helsinki and was reviewed and approved by the Ethics Committee of the Hospital District of Southwest Finland (ETMK:31/180/2011).

### MRI acquisition

The participants underwent an MRI scan solely for research purposes and without clinical indications. The scanning was performed at the Medical Imaging Centre of the Hospital District of Southwest Finland by an experienced radiographer, without anaesthesia, during natural sleep using the “feed and swaddle” procedure (Lehtola et al., 2019). We used a Siemens Magnetom Verio 3T scanner (Siemens Medical Solutions, Erlangen, Germany). The 60-minute protocol included a PD-T2-TSE (Dual-Echo Turbo Spin Echo) sequence with a Repetition Time (TR) of 12,070 ms and effective Echo Times (TE) of 13 ms and 102 ms (PD-weighted and T2-weighted images respectively), and a sagittal 3D T1-weighted MPRAGE sequence with 1.0 mm^3^ isotropic voxels, a TR of 1900 ms, a TE of 3.26 ms, and an inversion time (TI) of 900 ms. The total number of slices was 128 for both the T1- and T2-weighted images, and the images covered the whole brain. Sequence parameters were optimized so that “whisper” gradient mode could be used in the PD-T2-TSE and 3D T1-sequences to reduce acoustic noise during the scan. Single shell diffusion-weighted data was acquired with a standard twice-refocused Spin Echo-Echo Planar Imaging (SE-EPI) sequence (field of view (FOV) 208 mm; 64 slices; TR 9300 ms; TE 87 ms), with 2 mm^3^ isotropic resolution and a b-value of 1000 s/mm. There were in total 96 unique diffusion encoding directions in a three-part DTI sequence. Each part consisted of uniformly distributed 31, 32 or 33 directions and three b0 images (images without diffusion encoding) that were taken in the beginning, in the middle, and in the end of each scan (Merisaari et al., 2019, 2023).

All the brain images were assessed by a paediatric neuroradiologist for any incidental findings and, if the participant was found to have one, the infant and the parents were given a chance for a follow-up visit by a paediatric neurologist (Merisaari et al., 2023). Developmental status has thereafter been normal for all the participants, including those with incidental findings. The incidental findings were deemed not to affect brain anatomy / volume estimates of the participants in the current study. It is important to note that the encountered incidental findings have been found to be common and clinically insignificant in previous studies; see our recent article for more details (Kumpulainen et al., 2020). We have also covered the details of our scanning visits and tips for investigators in our review (Copeland et al., 2021).

### Template creation

#### Creation of population-specific FBN-125 structural templates

The images that were not suitable for data analysis (excessive number of artefacts) were excluded, leaving 125 / 180 successful structural MRI for template creation (69.4% success rate), which is comparably low and was due to technical issues with scanning that went unnoticed during data collection (Copeland et al., 2021). The MRIs that passed this quality control were used to construct a population-specific dual-contrast template (Figure 1 A). The T1 template was first created from T1 images and linearly registered to the MNI 152 template (Fonov et al., 2011). The average scaling from the native MRIs to the MNI 152 template was then computed, and the inverse was used to scale the MNI 152 template to the average size of our neonate population, which served as an initial target for construction of the population-specific template. The T2 images were linearly registered to the T1, and subsequently to the neonate template space with the transforms estimated from T1 scans.

**Figure 1.**
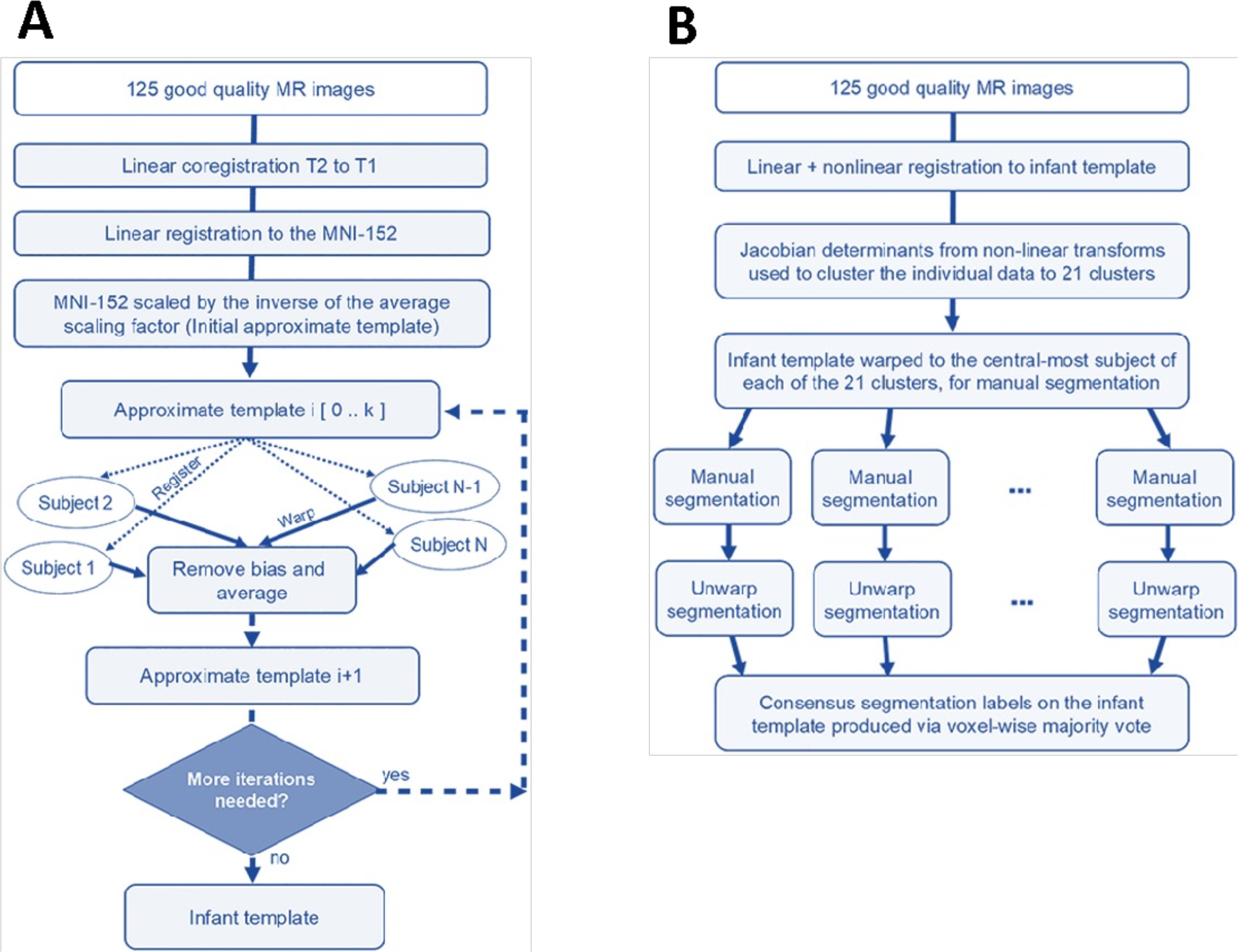
Summary of the workflow for template creation. A) Iterative construction of the infant template as described in Fonov et al. (2011). B) Labelling the infant template. The data were registered to the infant template, and then clustered based on the amount of distortion required to do that, into 21 clusters representing the morphological variability in the population. The template was then warped to the central-most subject of each cluster, providing 21 subtemplates for manual segmentation. After manual segmentation, the labels were then unwarped back to the base infant template and merged via voxel-wise majority vote to create the consensus labels. This figure is modified from Acosta et al. Cerebral Cortex, 2020 | reprinted with permission.

The template construction procedure is described in a prior article by Fonov et al. (2011) and is based on the work of Guimond et al. (2001); the method employs the principles of average model construction using elastic body deformations from Miller et al. (1997). It is an iterative procedure that, given a set of MRI volumes, builds a template which minimizes the mean squared intensity difference between the template and each subject’s MRI, and minimizes the magnitude of all deformations used to map the template to each subject’s MRI.

#### Creation of 21 subtemplates for manual segmentation

The non-linear transformations derived in the construction of the template were then used to cluster the subjects into 21 clusters from which we used the center-most subject as the basis to construct 21 targets for manual segmentation (Figure 1 B). As the basis for clustering, the Jacobian was computed for the non-linear transform mapping each subject to the template. The values in the Jacobian were extracted as a vector for each voxel within the template brain mask and clustered using an equal combination of cosine similarity and Euclidean distance with Ward’s clustering method (Ward Jr., 1963). We chose the number of clusters to be 21, which provided a good balance between reliable analysis procedures and the amount of work needed for manual labelling.

Then, within each of the 21 clusters, the sum-squared distance from each subject to each other subject was computed, and the subject with the minimum sum-squared distance was taken as the central-most subject of the cluster. The dual-contrast template constructed in the previous step was then warped to overlay the MRIs of these 21 subjects. These 21 subtemplates were then provided for manual segmentation without those doing that segmentation being made aware that these were, in fact, 21 versions / warped copies of the template. The demographics of the neonates whose brain images were determined to be one of the 21 cluster centroids are provided in Table 2.

**Table 2.**
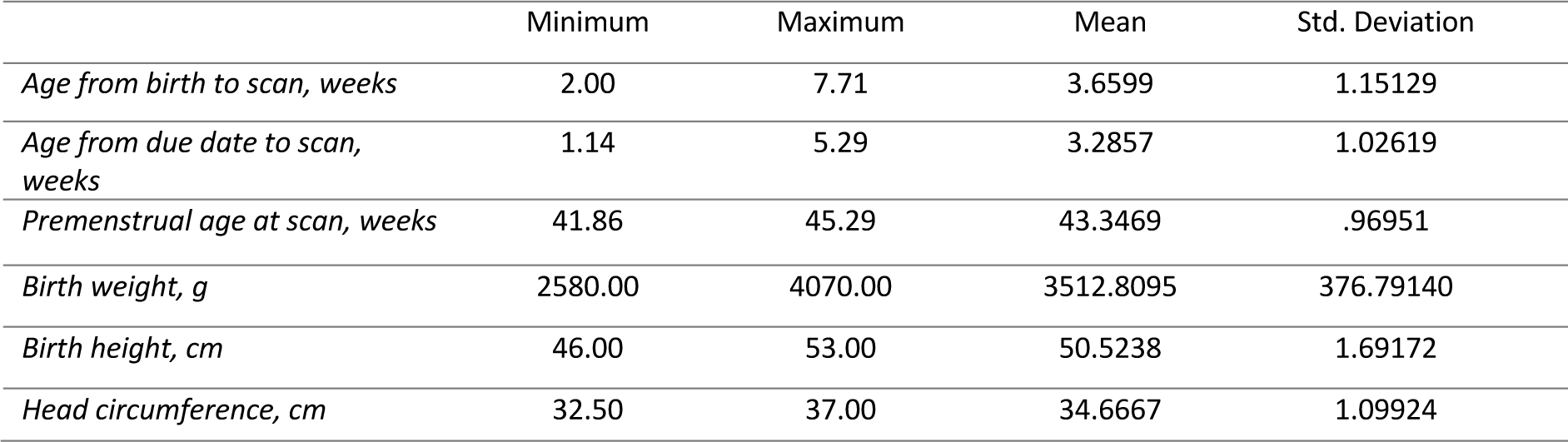
Demographics of the 21 neonates whose brain MRI scans were used as the basis to create the 21 subtemplates. These subtemplates were later used in the manual segmentation that yielded the segmented labels (7 male, 14 female).

|  | Minimum | Maximum | Mean | Std. Deviation |
| --- | --- | --- | --- | --- |
| *Age from birth to scan, weeks* | 2.00 | 7.71 | 3.6599 | 1.15129 |
| *Age from due date to scan, weeks* | 1.14 | 5.29 | 3.2857 | 1.02619 |
| *Premenstrual age at scan, weeks* | 41.86 | 45.29 | 43.3469 | .96951 |
| *Birth weight, g* | 2580.00 | 4070.00 | 3512.8095 | 376.79140 |
| *Birth height, cm* | 46.00 | 53.00 | 50.5238 | 1.69172 |
| *Head circumference, cm* | 32.50 | 37.00 | 34.6667 | 1.09924 |

#### Creation of FBN-125 DTI templates

Good quality b0 images were chosen manually, coregistered, averaged and moved in front of each 4D series. Brain masks were created based on the b0 volumes with the Brain Extraction Tool (BET; Smith, 2002) from FSL (FMRIB Software Library v 5.0.9; Jenkinson et al., 2012. DTIPrep software (Oguz et al., 2014) was used to inspect the quality of the data. Low quality diffusion images identified by DTIprep were discarded. The remaining images were then visually inspected following the automated quality control of DTIprep, and more directions were excluded as needed. We have found that after the quality control steps, datasets that have more than 20 diffusion encoding directions will yield reliable tensor estimates (Merisaari et al., 2019, 2023). Here all infants with at least 20 diffusion encoding directions were selected and we used all available participant’s data thereafter (N = 122). Eddy current and motion correction steps were conducted with FSL (Andersson & Sotiropoulos, 2016) and the b-vector matrix was rotated accordingly. A diffusion tensor model was fitted to each voxel included in the brain mask using the DTIFIT tool in FDT (FMRIB’s Diffusion Toolbox) of FSL using ordinary least squares (OLS) fit. Our DTI preprocessing steps have been provided in detail in our previous publications that also report good test-retest repeatability in between segments of the multi-part DTI sequences (Merisaari et al., 2019, 2023). The DTI template creation was carried out by rigidly registering the b0 images to the nonuniformity-corrected T1-weighted data and combining the transformations from b0-to- T1 and the T1-to- The FinnBrain Neonatal (FBN-125) template space for FA and MD maps (Acosta et al., 2020; Lewis et al., 2019). The registrations were carried out with ‘antsRegistration’. The FA and MD template images were then created by averaging the images with FSL’s fslmaths, part of FMRIB Software Library v6.0 (Jenkinson et al., 2012).

### Manual segmentation

#### The manual segmentation procedures and tools

Manual neonate brain segmentation is extremely labour intensive and requires considerable knowledge of the developmental characteristics of the various tissues. Full manual segmentation of the brain including cortical and subcortical grey matter, white matter and the CSF slice-by-slice is very time consuming. In our experience, working at 1 mm^3^ resolution this task takes around 1 month of full-time work. A higher 0.5 mm^3^ resolution of our subtemplates would have made this process even more labour intense, and we thus employed a model where we start the work from *initial estimates* for the gross tissue segmentation as outlined below. We were able to divide the work among research assistants whom we quickly, and successfully trained to perform the manual segmentations.

Another key thing that affected the workflow was the good initial tissue contrast in the created 21 subtemplates (due to averaging). Namely, the brain structures and their boundaries against neighbouring structures are relatively easy to detect. Manual segmentation is always prone to inter-rater and even intra-rater discrepancies, which may affect the statistical power in studies and lead to inaccurate estimates of outcome metrics. Here the use of 21 subtemplates to delineate the final segmentation on the FBN-125 template alleviates the final effect of minor errors and variability that stems from using multiple raters, and additionally allowed us to quantify the quality of segmentations.

We used teams of junior raters, supervised by senior investigators, to accomplish the work. For the subcortical grey matter nuclei, hippocampus, and amygdala, we started with one template jointly segmented for all subcortical structures by the primary rater NH and senior rater JJT (externally reviewed by JDL). The final subcortical segmentations of the 21 subtemplates were performed by three research assistants, supported by author NH on a regular basis, and all working under the supervision of JJT. The final labels on the 21 subtemplates were critically reviewed and corrected by JJT for consistency, and externally reviewed by JDL. The final labels are thus a consensus between two senior raters. The manual segmentation of amygdala, hippocampus and subcortical grey matter nuclei were done with Display software, part of MINC Tool Kit (https://bic-mni.github.io/).

For the cortical and gross anatomy segmentations, we made prior estimates of the structures that we manually corrected. The segmentations were performed by three research assistants. To aid the work, JJT prepared a detailed manual and video material showing model edits on each step. JJT also performed weekly quality control and checking all the segmentations as well as final check on all the images. As before, the images were externally reviewed by JDL. The gross anatomical segmentations were done with FSL tools and manual segmentation with fsleyes (McCarthy, 2021).

#### The manual segmentation of bilateral hippocampus and amygdala

We developed a detailed protocol for amygdala and hippocampus, which is provided in our prior article (Hashempour et al., 2019). Of important note, while we used identical procedures for amygdala and hippocampus segmentation, the change in image resolution from 1 mm^3^ to 0.5 mm^3^ enabled much more precision on the segmentations.

#### The manual segmentation of bilateral caudate nucleus

For the caudate, we decided to include ventral and dorsal caudate to the same label as there were no reliable landmarks for more fine-grained tracings (e.g. for nucleus accumbens), and the manual segmentation was based on prior work (Perlaki et al., 2017). We used the sagittal plane to trace the curvature of the caudate at the midline and then used the coronal plane as the main segmentation plane. After adjusting the contrast, the tissue borders were the main guides for labelling, and we used standard contrast range to assure systematic delineation of white matter and CSF boundaries. The key anatomical landmarks used for the tissue boundary detection were the lateral border of the lateral ventricles medially, white matter forming the external capsule and adjacent areas laterally, the anterior horns of the lateral ventricles and the capsula interna anteriorly, as well as the anterior commissure, putamen and globus pallidus for the anterior-inferior border.

#### The manual segmentation of bilateral putamen and globus pallidus

Once the caudate segmentation was ready, we segmented the putamen (mainly in the coronal plane). The anterior border was defined by the caudate head and otherwise the bilateral putamen were carefully traced with standard contrast settings to help systematic border delineation from the white matter while also keeping the claustrum separate from the tracings. The globus pallidus was segmented after the putamen starting from the posterior border and moving in circular tracings in all planes using a “lasso technique” where the outlines were first traced carefully in “easy-to-see” planes and the boundaries were then connected in other planes.

#### The manual segmentation of bilateral thalamus

The main approach for the thalamus segmentation was guided by prior work (Owens-Walton et al., 2019) and aided by the lasso technique (see above). The final and most challenging task is to assure that the tissue boundaries are refined and checked systematically in all planes to assure three-dimensional accuracy, which was checked and assured by the senior raters (all initial segmentations needed minor edits). The senior raters paid special attention to systematic separation of the corticospinal tract and frequently inferiorly spanning (pre)myelinated white matter and the inclusion of inferior parts of the thalamus that have different contrast features to the rest of the nucleus.

#### The manual segmentation of gross anatomical areas: cortical grey matter, white matter, deep grey matter, pons, and the cerebellum

To decrease the time needed for manual segmentations, we searched for the best initial segmentation from a selected range of software. Since none of the tried segmentations were perfect, the initial estimates for the 21 subtemplates were created with the FSL-VBM pipeline using the UNC neonate template grey matter probability mask to guide the segmentation (Douaud et al., 2007).

The manual correction for the *initial estimates* was performed in 21 steps as follows:

1. Erode the pre-estimates brain mask with fslmaths (creating eight versions with increasing eroding)
2. Mask the grey matter prior with a chosen “best fit” eroded brain mask to remove the mis-segmented GM outside the dural borders
3. Mask white matter prior with a chosen “best fit” eroded brain mask to remove the mis-segmented white matter outside the pial surface
4. Clean the midline from mislabelled white matter outside the pial surface
5. Clean the superior parietal areas from mislabelled GM and WM
6. Remove the mislabelled GM from the corticospinal tract and longitudinal fasciculus and fill them with the correct WM label
7. Fill in the central holes in the WM (near the subcortical grey matter nuclei)
8. Fill in the central holes in the subcortical GM that are mislabelled as CSF
9. Remove the mislabelled GM and WM from the inferior orbitofrontal regions
10. Segment the inferior temporal cortex as grey matter (fill in the holes in the cortex)
11. Remove the brainstem and cerebellum from the grey matter mask
12. Remove the subcortical grey from the previous image to yield cortical grey matter
13. Use fslmaths to calculate cortical grey matter, “deep grey matter” that included subcortical grey matter and adjacent myelinated white matter and brainstem-cerebellum (using the images from steps 11 and 12)
14. Segmentation of the cerebellum with major edits on the superior border
15. Fine segmentation of the cortical grey matter (thorough slice-wise visual inspection in all planes)
16. Fine joint segmentation of the brainstem and the cerebellum
17. Remove the brainstem from the joint segmentation
18. Use fslmaths to estimate separate the cerebellum, brainstem and cerebral white matter (using all estimated parts obtained at this stage and step 13)
19. Review all segmentations separately and edit if needed
20. Use fsleyes to smooth and fslmaths to threshold and binarise the reviewed segmentations. This step alleviates the occasional “ragged edges” that are common after manual edits
21. Make intracranial volume (ICV) and CSF masks using the brain mask and images obtained from step 19
22. Fine segmentation of the CSF that included segmentation of the internal CSF within the lateral ventricles

All steps were followed by visual quality control, i.e. moving across the edited areas with variable speed and in all viewing planes. We identified brain regions that are typically challenging for automated pipelines, and these were reviewed with special care:

1. Breaks in the inferior (thin) cortex – temporal and occipital lobes
2. Overestimation of very thin cortical strips in the parietal areas towards the dura / skull
3. Areas with no visible csf, grey matter fusion with the skull / dura in superior posterior parts
4. Overlap between cerebellum and occipital lobes and mixing with the transverse sinus
5. Cingulate and corpus callosum mis-segmentation
6. Errors in the deep grey segmentation due variability in the anatomy / myelination
7. Amygdala and hippocampus segmentation errors (frequently to all directions!)

The final output of the manual segmentation were binary labels for cortex, white matter, internal and external CSF (the labels were later combined), brainstem, cerebellum, and a “deep-grey” segmentation that intentionally covered the subcortical grey matter and the myelinated portions of white matter surrounding the nuclei. The previously created subcortical areas were subtracted from this label and the remaining voxels were added to the binary white matter mask.

### The creation of atlas labels from manual segmentations

After manual segmentation, the labels were unwarped back to the FBN-125 space and merged via voxel-wise majority vote to create the consensus labels, and the labels were assured to be symmetric and complete through visual inspection.

Finally, we used the symmetric labels to create several atlases: 1) gross tissue labels for grey matter, white matter and CSF; 2) symmetric labels for grey matter, white matter, CSF, brainstem and cerebellum as well as labels of the bilateral amygdala, hippocampus, caudate, putamen, globus pallidus and thalamus; and 3) corresponding asymmetric labels for left and right hemispheres with FreeSurfer look up table labels. For the creation of the asymmetric labels, we defined a right hemispheric binary mask to aid the separation of the hemispheres. The labels were created from symmetric labels with ‘fslmaths’ from the FMRIB Software Library v6.0 (Jenkinson et al., 2012). The FreeSurfer labels were obtained from: (https://surfer.nmr.mgh.harvard.edu/fswiki/LabelsClutsAnnotationFiles).

We calculated the generalized conformity index (GCI) for all structures to quantify the agreement across the atlas labels. Here, the GCI quantified the spatial overlap among the manually defined atlas labels. GCI is a generalization of the Jaccard score so that for two raters the GCI equals the Jaccard score, GCI = Vol(A1 ∩ A2)/Vol(A1 ∪ A2). We quantified the GCI across the 21 manual segmentations by including segmentation j, its volume Vol(Aj), and Σ pairs (i>j) the summation over all combinations of unique pairs of labels, and defined GCI as:

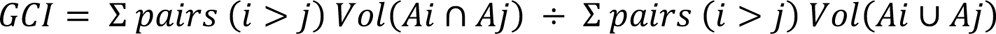

We first binarized each manually created label and then added all unique pairs of these binarized labels so that all voxels with a value > 0 as their union, and all voxels with value 2 as their intersection. We then used ANTs ‘LabelGeometryMeasures’ to calculate the size of both the union and intersection and used those values in the formula for GCI (Kouwenhoven et al., 2009; Visser et al., 2019). Since the FBN-125 template is symmetric, we reported an average of bilateral labels.

### Benchmarking transfer of statistical maps of functional MRI activations from neonatal to adult MNI space

We created standard coregistration files from the FBN-125 neonate template to the adult MNI space and make them freely available with the templates and atlases. For these transforms we estimated a transform from the adult MNI space template to the FBN-125 neonate template to prevent the effects the (minor) differences in cortical anatomy to the final transforms. We used ‘antsRegistrationSyNQuick.sh’ and ‘antsApplyTransforms’ available from ANTs software for all coregistrations (Avants et al., 2011; Tustison & Avants, 2013).

The templates used for spatial normalization in neonatal studies vary from using an MNI template (Wild et al., 2017) or standard Talairach space (Biagi et al., 2015) for adults and for infants a study-specific template (Dehaene-Lambertz et al., 2002) or off-the-shelf atlas, (Goksan et al., 2015; Wild et al., 2017) such as the UNC infant template (Shi et al., 2011; Wild et al., 2017). We chose the UNC-0-1-2-years neonate template as the model template for coregistrations as it was identified as is the most used off-the-shelf atlas used for infants (Dufford et al., 2022).

To test the utility of transferring statistical maps obtained in neonatal functional MRI (fMRI) from neonatal template space to adult MNI space, we used results from our recent fMRI study (Mariani Wigley et al., 2023). We first estimated transforms from UNC neonate template to FBN-125 template space. We then transformed statistical maps to adult MNI space by concatenating the warps from UNC – to FBN-125 – to MNI template space.

## Results

### High-resolution multimodal neonatal brain templates for structural and diffusion MRI, and accompanying atlases

FBN-125 templates entail a set of multi-contrast template volumes of T1 - and T2 – weighted templates (Figure 2A and 2B), corresponding DTI tensor templates of FA and MD average maps (Figure 2C and D) and accompanying atlases with gross anatomical (Figure 3A), symmetric (Figure 3B), and asymmetric labels (Figure 3C).

**Figure 2.**
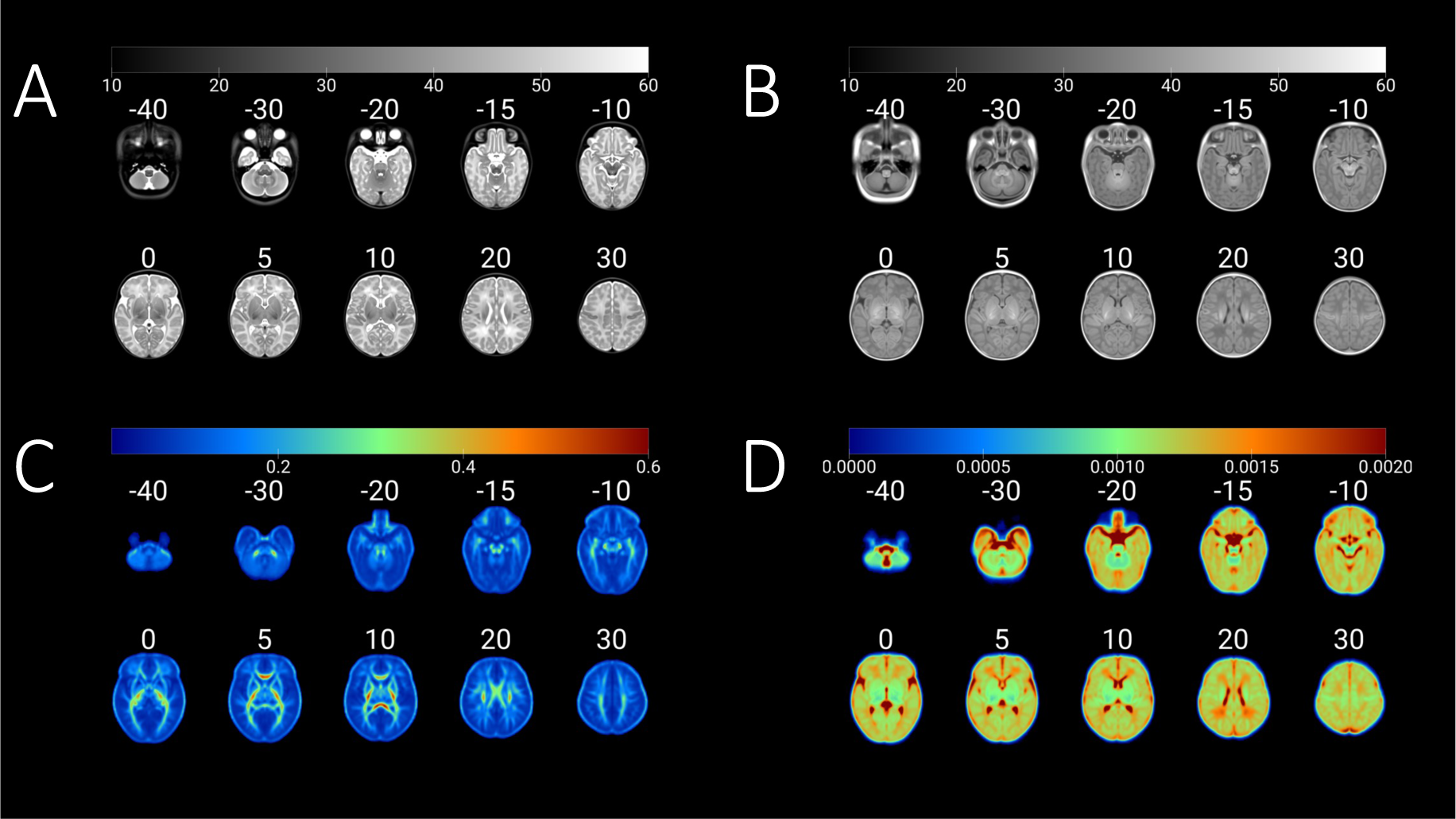
The FinnBrain Neonate FBN-125 templates for A) T2-weighted, B) T1-weighted, C) DTI-derived fractional anisotropy, and D) mean diffusivity. The grey colour scales depict intensity for A and B, and DTI tensor scalar values for C (untiless) and D (mm^2^/s). Each axial slice has been tagged with a z coordinate of the adult MNI template space (in MRIcroGL software).

**Figure 3.**
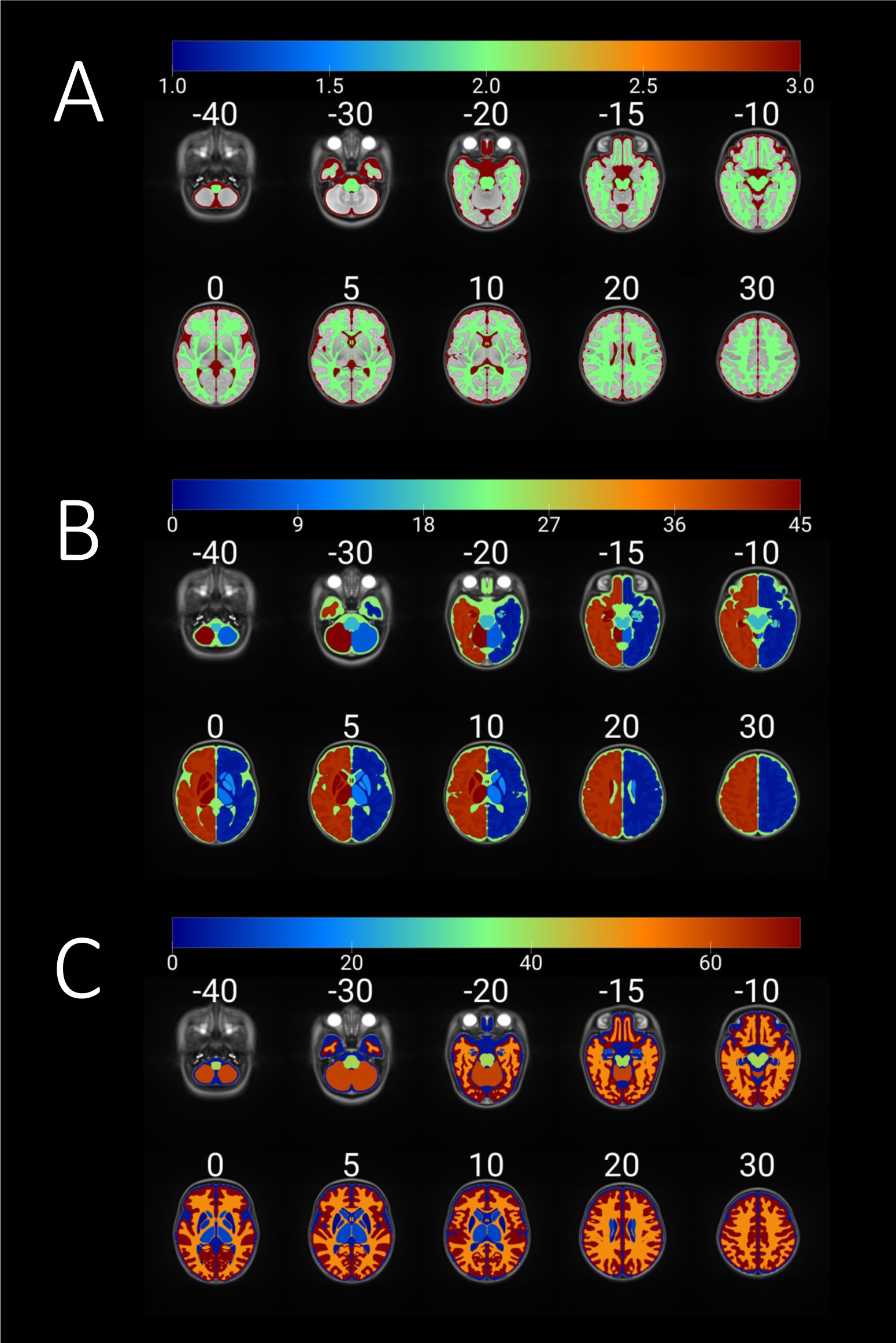
The anatomical labels for FBN-125 templates include A) gross anatomical labels of grey and white matter as well as CSF, B) asymmetric labels in FreeSurfer lookup table (LUT) compliant form, and C) symmetric labels. Colour scales depict anatomical label numbers. Each axial slice has been tagged with a z coordinate of the adult MNI template space (in MRIcroGL software).

### Consistency of manual labels used to create the atlases

The agreement of the manual segmentations for the 21 subtemplates was good for all structures: GCI ranged between 0.71 – 0.86 (Table 3). GCI scores of 0.7–1.0 are regarded as excellent (Kouwenhoven et al., 2009; Visser et al., 2019).

**Table 3.** Manual segmentation accuracies measured with Generalized Conformity Index (CGI).

| Region of interest | GCI |
| --- | --- |
| Caudate | 0.81 |
| Putamen | 0.81 |
| Globus pallidus | 0.71 |
| Thalamus | 0.86 |
| Hippocampus | 0.75 |
| Amygdala | 0.71 |
| White matter | 0.83 |
| Cortex | 0.80 |
| Cerebrospinal fluid | 0.80 |
| Brainstem | 0.86 |
| Cerebellum | 0.92 |

### Novel means to transform neonatal functional MRI results to adult MNI standard space

We estimated standard transforms from the FBN-125 template to the adult MNI template space. We then transformed statistical maps obtained in our prior study reporting brain activations to social touch in neonates (Mariani Wigley et al., 2023) to adult MNI space. The registrations were accurate (Figure 4) and worked equally well for unthresholded T maps.

**Figure 4.**
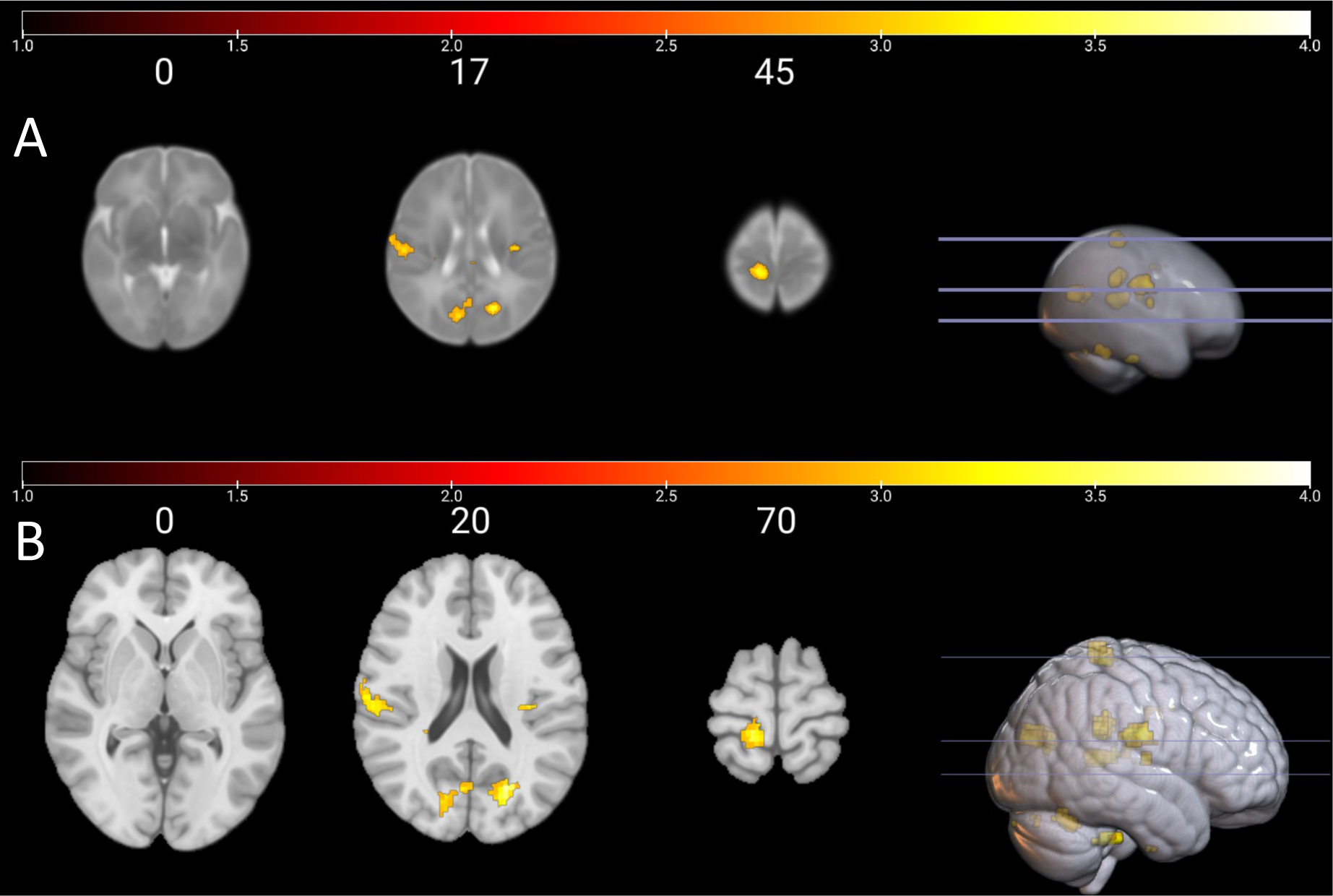
Main effect of brushing vs. rest conditions in neonates A) in the UNC neonate atlas template space as in Mariani Wigley et al., 2023, and B) in the adult MNI space (mni_icbm152_t1_tal_nlin_sym_09a) after transforms to FBN-125 neonate template space and using the standard transforms from FBN-125 neonate to adult MNI space. The colourbars visualise T values from thresholded cluster p < 0.005, FDR corrected for multiple comparisons (N = 18), see Mariani Wigley et al., 2023 for more information. Adult MNI space z coordinates appear on top of each axial slice. Note the different choice of coordinates in UNC/MNI template spaces to visualise the same regions of interest from the contrast in both images.

## Discussion

We created a novel set of neonatal templates and atlases with the key novel contribution of enabling standard spatial transformations between neonatal and adult MNI space. This is potentially an important step in standardizing the use of template spaces, which according to a recent review is much needed (Dufford et al., 2022), and also enabling comparisons between neonates and adults. Second, a related contribution is that we created multimodal templates for structural and diffusion MRI, which are rare in the field (Table 1, included as supplement 1 here). We make the FBN-125 neonate templates, atlases and adult MNI coregistration files publicly available for the scientific community (see Data Sharing).

### Potential for better comparability for neonatal MRI studies

When reporting findings from a neuroimaging study, it is important to specify which template and coordinate space was used for spatial normalization so that data collected using different methods can be compared across studies (Poldrack et al., 2008). For adults, the MNI 152 template is the most frequently used standardized template space for spatial normalization (Mazziotta et al., 2001a, Mazziotta et al., 2001b, Mazziotta et al., 1995). However, infant neuroimaging research predominantly processes infant data in single subject space due to a lack of a standardized template (Dufford et al., 2022). Usage of off-the-shelf infant templates followed by study-specific templates were most common in studies using fMRI (Dufford et al., 2022). Specifically, in term-born populations, 81 studies used off-the-shelf atlases, 29 studies used a study-specific common space, and 16 studies used a single subject space, indicating strong preferences for off-the-shelf atlases. The most commonly used template / atlas across modalities were the UNC infant atlases (24%) and the JHU neonate atlases (13%) (Dufford et al., 2022).

In the case of task-based fMRI studies, reporting the coordinates of neural activity related to a specific task is useful for comparability and replication across studies. However, only some task fMRI studies with infant populations report the corresponding MNI coordinates of the activated brain areas (Ellis et al., 2021a; Ellis et al., 2021b; Graham et al., 2013). For instance, whilst some studies reported the region of interest (ROI) coordinates from an off-the-shelf infant template (Kuklisova-Murgasova et al., 2011; Williams et al., 2015), some report the coordinates of peak activity in Talairach space (Blasi et al., 2011). Further, in task fMRI studies comparing infant and adult samples, MNIspace coordinates are reported for the adult participants, whilst the infant coordinates are reported in correspondence to an off-the-shelf infant atlas (Goksan et al., 2015). Some studies have reported the locations so that coordinates from the adult sample are given in Talairach space, and coordinates in millimetre points from the anterior commissure point for the infant sample (Biagi et al., 2015). Many studies have opted to report the results in the predetermined ROIs (Allievi et al., 2016; Anderson et al., 2001; Baldoli et al., 2015; Deen et al., 2017; Dehaene-Lambertz et al., 2002; Lee et al., 2012; Perani et al., 2010; Wild et al., 2017), but it is clear that at the moment the benefits of standard space and coordinates cannot be fully established in neonatal and infant MRI studies.

The FBN-125 templates could provide a standard waypoint template for all existing and future studies to enable investigators to compare locations of their activations through adult MNI coordinates (https://neurosynth.org/), report standard MNI coordinates that enable meta-analyses (https://www.brainmap.org/ale/), and to store unthresholded T maps for later use (https://neurovault.org/). On a related note, recent advances in available longitudinal atlases spanning ages from gestation to the neonatal period (Serag et al., 2012) as well as from birth to age two years (Ahmad et al., 2023) can be integrated with our templates through serial registration across the longitudinal template series to neonatal template, registration to FBN-125 and standard transform to the MNI space.

### The typical features of the neonatal MRI, limitations, and future directions

The neonate brain is roughly one third of the adult brain, which makes the proportional resolution worse by a factor of 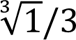 - e.g. 1 mm^3^ resolution in the neonate brain is equivalent to 1.5 mm^3^ in adult image. Overall, this makes the partial volume issues more pronounced. Second, the infant brain morphology usually has a lot more variance than in older ages (Li et al., 2019), e.g. the bones of the neonatal skull are not fused and there may be marked left–right asymmetries, flattening in either anterior–posterior or superior–inferior direction or even bulging of the brain out of the superior foramen – all this reflecting perfectly normal anatomy. Minor birth-related haemorrhages, incidental findings, are also important to consider (Kumpulainen et al., 2020), although they can often be dealt with via corrections in brain masks and selected exclusions of study participants. We performed the manual segmentations on the warped copies of the average template, which made the manual segmentation easier due to relatively “sharp” tissue borders. The initial averaging in template creation enabled both an increase in signal-to-noise ratio and up-sampling the resolution from the initial scan resolution (here from 1.0 mm^3^ to 0.5 mm^3^). Even small structures such as the claustrum are visible in the templates, and the images could be used to manually label additional, smaller structures in the future.

It is imperative to note that the infant brain tissue contrast changes in at least three different phases (Li et al., 2019): “(1) the infantile phase (≤3 months), in which the grey matter shows a relatively higher signal intensity than the white matter in T1w images, and the tissue contrast in T2w images is better than in T1w images; (2) the isointense phase (5–9 months), in which the signal intensity of the white matteer is increasing during the development due to the myelination and maturation process; in this phase, grey matter and white matter have the lowest signal differentiation in both T1w and T2w images; (3) the early adult-like phase (≥12 months), where the grey matter intensity is much lower than the white matter intensity in T1w images, largely similar to the tissue contrast pattern in adult T1w images.” Our atlas has been built from a Finnish (Scandinavian Caucasian) term-born population scanned at the gestation-corrected age of 1–5 weeks (age from birth 2–7 weeks) and is best suited for analyses on neonatal / early infancy data. Unfortunately, we have only a small number of participants with a follow-up scan after their neonatal scan, and we are thus not able to contribute to longitudinal atlas development across infancy. Consequently, our templates may not fit the needs of studies carried out in preterm populations that have also used a mixed set of templates (Allievi et al., 2016; Anderson et al., 2001; Lee et al., 2012). The joint efforts of large-scale projects such as the Developing Human Connectome Project, Baby Human Connectome Project, and Healthy Brain Child Development will provide high quality data and related software to support 4D atlas development from infancy to early childhood and beyond (Ahmad et al., 2023; Serag et al., 2012). We propose that future studies that introduce new templates and atlases would include standard transforms to the adult MNI space as was done in the current study.

## Conclusions

Neonatal brain segmentation remains a key challenge for developmental neuroscience. Advances in the field may rely on producing better templates, atlases, and segmentation tools. We contribute to this endeavour here by creating and sharing our FBN-125 neonatal templates, atlases, and standard registrations between neonatal and adult standard spaces. The created labels are amenable to coregistration to diffusion or functional scans, e.g. for tractography and seed-based connectivity analyses. Finally, other groups can contribute to manual labelling of additional structures, hopefully in time producing increasingly detailed atlas labelling.

## Data sharing

We plan to make the FBN-125 templates and atlases publicly available for the scientific community, and also provide the standard coregistration files between the FBN-125 and adult MNI spaces (details will be made available later following peer reviewed publication). We are happy to share insights in formal collaboration and interested investigators are encouraged to contact the corresponding author JJT. Raw and derived data sharing is possible via formal material / data sharing agreements that can be made by contacting the FinnBrain administration. The up-to-date contact information can be found from (https://sites.utu.fi/finnbrain/en/contact/).

## Acknowledgements

We would like to warmly thank all FinnBrain families that participated in the study. We would also like to thank the research team that had a supportive role in the current study: Satu Lehtola for her help in data collection, Maria Lavonius for her help in recruiting the participants, Jani Saunavaara for implementing the MRI sequences, Riitta Parkkola for reviewing the MR images for incidental findings.

## Author contributions

- **Jetro J. Tuulari,** conceptualization, creation of manual segmentation guidelines, supervising the manual segmentation work, creation of the DTI templates, creating the atlas labels, validating the standard coregistrations, drafting the initial version of the manuscript, leading the manuscript writing.
- **Aylin Rosberg,** performing manual segmentation, drafting the initial version of the manuscript.
- **Elmo P. Pulli,** drafting the initial version of the manuscript, providing feedback to early versions of the manuscript.
- **Niloofar Hashempour,** performing manual segmentation, creation of manual segmentation guidelines, co-supervising the manual segmentation work of the subcortical structures, drafting the initial version of the manuscript.
- **Elena Ukharova,** performing manual segmentation, drafting the initial version of the manuscript.
- **Kristian Lidauer,** performing manual segmentation, drafting the initial version of the manuscript.
- **Ashmeet Jolly,** drafting the initial version of the manuscript.
- **Silja Luotonen,** drafting the initial version of the manuscript.
- **Hilyatushalihah K. Audah,** drafting the initial version of the manuscript.
- **Elena Vartiainen,** drafting the initial version of the manuscript.
- **Wajiha Bano,** drafting the initial version of the manuscript.
- **Ilkka Suuronen,** drafting the initial version of the manuscript.
- **Isabella Wigley,** validating the standard coregistrations.
- **Vladimir S. Fonov,** developed the tools used in the template creation and segmentation of the data
- **D. Louis Collins,** developed the tools used in the template creation and segmentation of the data.
- **Harri Merisaari,** creation of the DTI templates, drafting the initial version of the manuscript.
- **Linnea Karlsson,** planned and funded the MRI measurements, established the FinnBrain Birth Cohort and built the infrastructure for carrying out the study.
- **Hasse Karlsson,** planned and funded the MRI measurements, established the FinnBrain Birth Cohort and built the infrastructure for carrying out the study.
- **John D. Lewis,** conceptualization, creation of the MRI sequences, creation of manual segmentation guidelines, supervising the manual segmentation work, performing the segmentation of the ‘base template’ creating the atlas labels, validating the standard coregistrations, drafting the initial version of the manuscript, leading the manuscript writing.

**All authors** participated in writing the manuscript and accepted the final version.

