## supplement 1 for "The FinnBrain Multimodal Neonatal Template and Atlas Collection: T1, T2, and DTI brain templates, and accompanying cortical and subcortical atlases"

**Table 1.** Review of available neonatal and infant templates and atlases. N = number of participants, M = MRI modalities, L = number of labels in the parcellated atlas, K = number of time points, R = resolution, A = ages of the infants, S = related segmentation procedures/software.

| Study | N | M | L | K | R | A | S |
|---|---|---|---|---|---|---|---|
| **Akiyama Atlas (Akiyama et al., 2013)**<br><br>https://ilabs.uw.edu/6-m-templates-atlas/ | 60 (29 males, 31 females) | Structural: T1 – acquired from two sites; 1.5 T scanner (N = 27: 14 males, 13 females; only sagittal images); 3T scanner (N = 33: 15 males, 18 females) | 90 (AAL; Anatomical Automatic Labeling) | 1 | 1.5T scanner-0.94×0.94×1 $mm^3$; 3T scanner-1×1×1 $mm^3$ | 6 months | Extracted 6-month-old average brain was segmented into brain tissue and CSF using FSL FMRIB's Automated Segmentation Tool (FAST); ANTS. Template construction using tools from the Medical Image NetCDF (http://www.bic.mni.mcgill.ca/ServicesSoftware).<br><br>https://pubmed.ncbi.nlm.nih.gov/24069234/ |
| **Altaye Atlas (Altaye et al., 2008)**<br><br>http://www.irc.cchmc.org/software | 77 (31 males, 46 females) | T1 | 3 (GM, WM, CSF) | 1 | 1×1×1 $mm^3$ | 9-15 months | Dual strategy segmentation approach using SPM5:<br>i) unified segmentation based on *a priori* adult segmentations<br>ii) current voxel-intensity approach based on a Gaussian mixture model. The final tissue probabilities are estimated without tissue priors. Both strategies apply an HMRF model (Cuadra et al., 2005) during template construction. |

| | | | | | | | |
|---|---|---|---|---|---|---|---|
| | | | | | | | https://pubmed.ncbi.nlm.nih.gov/18761410/ |
| **EBDS Neonatal DTI Atlas (Short et al., 2022)**<br><br>https://www.nitrc.org/projects/uncebds_neodti | Newborns 144 (68 males, 76 females); 1-year 170 (95 males, 75 females); 2-years 171 (88 males, 83 females). | DTI | A total of 47 tract segments were obtained for the neonatal DTI atlas (see Lee et al., 2015). | 3 | Slice thickness 2 mm and in-plane resolution = 2mm× 2mm | Newborn, 1 and 2 years | Fiber-tract-based analysis closely follows UNC Utah NAMIC DTI analysis framework (Verde et al., 2013, 2014) https://www.nitrc.org/projects/namicdtifiber/DTIAtlasBuilder).<br><br>Atlas construction using DTIAtlasBuilder (https://www.nitrc.org/projects/dtiatlasbuilder)<br><br>https://pubmed.ncbi.nlm.nih.gov/35401073/ |
| **Edinburgh Neonatal Atlas (ENA33) (Blesa et al., 2016)**<br><br>https://git.ecdf.ed.ac.uk/jbrl/ena/tree/00fe2c0ae25f326338369175643356acf272f780 | 33 (NN male/female) | Structural T1, T2, DTI | 107 (SRI24/TZO) | 1 | T1: 1×1×1 $mm^3$<br><br>T2: 0.9×0.9×0.9 $mm^3$<br><br>DTI: 2mm | Term-born, 37-41 PMW | Tissue segmentation used nonlinear registration to the closest age-matched T1w template from the 4D atlas using Free-Form Deformation implemented in NiftyReg, expectation-maximization (EM) algorithm used to classify each voxel into tissue. |

| | | | | | | | |
|---|---|---|---|---|---|---|---|
| | | | | | slice thickness | | The neonatal brain was parcellated into the SRI24/TZO adult brain atlas (Rohlfing et al., 2009) using the LISA method (Serag et al., 2012) to model the anatomical differences between adult and neonatal brains. Template and atlas were constructed using the Symmetric Group Normalization (SyGN) method.<br><br>https://pubmed.ncbi.nlm.nih.gov/27242423/ |
| **Imperial ALBERTs (Gousias et al., 2012)**<br><br>https://brain-development.org/brain-atlases/neonatal-brain-atlases/neonatal-brain-atlas-gousias/ | 20: 15 preterm (7 males, 8 females); 5 term-born (3 males, 2 females) | Structural T1 and T2 | 50 | 1 | T1: 0.82×0.82×1.6 $mm^3$ (resliced to 0.82×0.82×0.82 $mm^3$)<br><br>T2:0.86×0.86×2.0 $mm^3$ | Preterm: median 40 weeks (range 37-43 weeks); term-born: median 41 weeks (range 39-45 weeks) | Manual segmentation of the whole brain into 50 regions based on previous protocols (Ahsan et al., 2007; Gousias et al., 2008; Hammers et al., 2003, 2007) using macroanatomical landmarks. Each voxel was labelled as belonging to one ROI, resulting in a label-based encephalic ROI template (ALBERT).<br><br>https://pubmed.ncbi.nlm.nih.gov/22713673/ |

| | | | | | | | |
|---|---|---|---|---|---|---|---|
| **Imperial Neonatal Atlas (Kuklisova-Murgasova et al., 2011)**<br><br>https://brain-development.org/brain-atlases/neonatal-brain-atlases/neonatal-brain-atlas-murgasova/ | 142 (70 males, 72 females) | Structural T2 | 6 (cortex, white matter, subcortical grey matter, brainstem, cerebellum and cerebrospinal fluid) | 15 (gestational weeks 29-44) | 0.86 × 0.86 × 1 $mm^3$ | In the neonatal population, t = 36.6 gestational weeks (range 28.6-47.7 gwk, SD ± 4.9 gwk), | Brain segmentation with an intensity-based approach similar to the work of (Xue et al., 2007), using atlas-based segmentations based on manual delineations of deep grey matter, brainstem, cerebellum and darker regions of white matter. Kernel-based regression method was used for template and age-specific 4D probabilistic atlas creation (Davis et al., 2010; Ericsson et al., 2008).<br><br>https://pubmed.ncbi.nlm.nih.gov/20969966/ |
| **Imperial Pediatric Atlas (Gousias et al., 2008)**<br><br>https://brain-development.org/brain-atlases/pediatric-brain-atlases/pediatric-brain-atlas-gousias/ | 33 (17 males, 16 females) | Structural: T1 (only images acquired in the sagittal plane) | 83 | 1 | 1.04 × 1.04 × 1.04 $mm^3$ | 1, 2 years (mean 24.8 months, median 24.1 months (range 21.4–34.4 months) | Automatic segmentation of pediatric brains using an algorithm that was based on manual segmentation of 30 adult brains that resulted in 30 adult atlases labelling 83 anatomical structures. Final segmentation combined the 30 adult atlases using decision fusion.<br><br>https://pubmed.ncbi.nlm.nih.gov/18234511/ |

| | | | | | | | |
|---|---|---|---|---|---|---|---|
| **Imperial Spatio-Temporal Atlas (Serag et al., 2012)**<br><br>https://brain-development.org/brain-atlases/neonatal-brain-atlases/neonatal-brain-atlas-serag/ | 204 neonatal (NN male/female) | Structural T1 and T2 | 6 (cortex, white matter, deep grey matter, brainstem, cerebellum and cerebrospinal fluid); 5 for fetuses (cortex, left and right hemispheres, ventricles, and cerebrospinal fluid) | 9 (28, 30, 32, 34, 36, 38, 40, 42, 44 PMW) | T1: 0.82×1.03×1.6 $mm^3$ T2: 1.15×1.18×2 $mm^3$ | M = 37.3 weeks (range 26.7-44.3 weeks, SD ±4.8 weeks) | Procedure following Imperial Neonatal Atlas (Kuklisova-Murgasova et al., 2011).<br><br>https://pubmed.ncbi.nlm.nih.gov/21985910/ |
| **Infant FreeSurfer Atlases (de Macedo Rodrigues, 2015)** | 23 (NN male/female) | Structural T1 | 32+14 | 26 | 1×1×1 | 0-2 years | Manual segmentation<br><br>https://pubmed.ncbi.nlm.nih.gov/25741260/ |
| **INSERM Atlas (Dehaene-Lambertz et al., 2002)** | 20 (6 males, 24 females) | T2 | 13 | 1 | 0.391×0.391×4 $mm^3$ (resampled at 3.1×3.1×4 $mm^3$) | 3 months | The template was constructed using manual alignment of the AC-PC commissures for two participants using SPM99 and Anatomist.<br><br>https://pubmed.ncbi.nlm.nih.gov/12471265/ |

| | | | | | | | |
|---|---|---|---|---|---|---|---|
| **JHU-neonate-linear and JHU-neonate-nonlinear-atlases (Oishi et al., 2011)**<br><br>https://cmrm.med.jhmi.edu/cmrm/Data_neonate_atlas/atlas_neonate.htm | 20 (15 males, 10 females) | Structural T1, (N = 14), T2 and DTI (N = 20) | 122 | 1 | T1, T2: 1×1×1 $mm^3$<br>DTI: 0.6×0.6×0.6 $mm^3$ | Term-born, 0-4 days after birth | Manual segmentation procedure follows the adult JHU-MNI parcellation map.<br><br>Each T2w image was aligned to a common AC-PC line based on the averaged brain size and shape. This was applied to all collected modalities creating the JHU neonate-linear atlas.<br><br>Co-registered DTI images were nonlinearly normalized to the JHU-neonate-SS atlas with a dual-channel LDDMM (Ceritoglu et al., 2009; Miller et al., 1997) using DiffeoMap (see (Oishi et al., 2009). The resulting nonlinear transformation matrices were applied to the co-registered T1w and T2w images creating the JHU-neonate-nonlinear-atlases.<br><br>https://pubmed.ncbi.nlm.nih.gov/21276861/ |
| **JHU-neonate-SS Atlas (Oishi et al., 2011)**<br><br>https://cmrm.med.jhmi.edu/cmrm/Data_n | 1 | Structural T1, T2 | 122 | 1 | T1, T2: 1×1×1 $mm^3$ | Term-born, 2 days after birth | Manual segmentation procedure follows the adult JHU-MNI parcellation map using ROIEditor |

| | | | | | | | |
|---|---|---|---|---|---|---|---|
| eonate_atlas/atlas_neonate.htm | | | | | | | (www.Mristudio.org) to inspect all three slice orientations.<br><br>One subject with the best fitting brain shape to the JHU neonate-linear-atlas was linearly normalized to the T2w image of the JHU-neonate-linear. The resulting transformation matrix was applied to the other co-registered DTI and T1w images creating the JHU-neonate-SS.<br><br>https://pubmed.ncbi.nlm.nih.gov/21276861/ |
| **M-CRIB Atlas (Alexander et al., 2017)**<br><br>https://github.com/DevelopmentalImagingMCRI/M-CRIB_atlas<br><br>M-CRIB 2.0<br><br>https://pubmed.ncbi.nlm.nih.gov/30804737/ | 10 (6 males, 4 females) | Structural T2 | 100 (cortical and subcortical labels matching the Desikan-Killiany parcellation (Desikan et al., 2006) | 1 | 0.63×0.63×0.63 $mm^3$ | Term-born, age at scan 40.29–43.00 gwk, M=41.71 | MANTiS for automatic tissue classification, then manual cleaning of tissue segmentation; all parcellation of high-resolution T2w images using ITK-SNAP (Yushkevich et al., 2006) and manual parcellation for 100 different regions.<br><br>Linear and nonlinear T1w and T2w templates were constructed using ANTS V2.1.<br><br>https://pubmed.ncbi.nlm.nih.gov/27725314/ |

| | | | | | | | |
|---|---|---|---|---|---|---|---|
| **MRICloud neonate multi-atlas repository (Otsuka et al., 2019)** | 7 (3 males, 4 females) | Structural T1 | 30 | 7 | 1×1×1 mm$^3$ | Term- and preterm-born, 38-42 PMW | Automatic parcellation using MALF integrated with segmentation tools in MRICloud (https://mricloud.org/). https://pubmed.ncbi.nlm.nih.gov/31037800/ |
| **Multi-structural Neonatal Brain Atlas (Makropoulos et al. 2016)**<br><br>(http://lbam.med.jhmi.edu/) | 338 (298 preterm; NN male/female) | T2 | 50 (WM, cortical and subcortical structures); becomes 82 in the atlas after further subdividing WM and cortical GM parts. | 5 (28, 32, 36, 40, 44 PMA) | Slice thickness 2 mm acquired with an overlap of 1 mm; in-plane resolution 0.86 × 0.86 mm | Median 39 (27-44) PMW | Segmentation protocol following (Makropoulos et al., 2014); Expectation-maximization algorithm; image intensity modelled with Gaussian Mixture Model.<br><br>A 4D spatio-temporal structural atlas of the brain built from the 82 cortical and subcortical segmentation averages.<br><br>https://pubmed.ncbi.nlm.nih.gov/26499811/ |
| **NIHPD Objective 2 Atlases (Fonov et al. 2009)**<br><br>http://www.bic.mni.mcgill.ca/ServicesAtlases/NIHPD-obj2 | 108 (NN male/female) | T1 | No anatomical parcellation provided | 11 | 1×1×3 mm$^3$ | 0-2, 2-5, 5-8, 8-11, 11-14, 14-17, 17-21, 21-27, 27-33, 33-44, 44-60 months | A suite of software developed by the Montreal Neurological Institute (MNI) was used.<br><br>*PMID N/A*<br>https://www-sciencedirect-com.ezproxy.utu.fi/science/article/pii/S1053811909708845 |
| **Singapore Atlas (Broekman et al. 2014)** | 93 (44 males, 49 females) | T2 and DTI | 2 (FA and DTI colour map) | 2 (*in utero* and shortly | DTI: 3.0 mm thickness | M = 9.9 days (SD = 2.3) | DTI Atlas was created using the unbiased diffeomorphic atlas generation algorithm (Joshi et al., 2004). FA image aligned to JHU- |

| | | | | | | | |
|---|---|---|---|---|---|---|---|
| http://www.bioeng.nus.edu | | | | after birth) | (axial slices) | | neonate-SS DTI atlas (Oishi et al., 2011); Voxel-based analysis using SPM8. https://pubmed.ncbi.nlm.nih.gov/25535959/ |
| **UNC-CH Longitudinal Infant Atlas (Kim et al. 2017)** | 28 (NN male/female) | DW | No anatomical parcellation provided | 2 (6 and 12 months) | 2×2×2 $mm^3$ | Term-born, age not reported | All DW images were processed using FSL. DW atlases were constructed by fusing diffusion-weighted images across time points and space in a patch-wise way using sparse representation with a graph constraint that promotes spatiotemporal consistency. https://pubmed.ncbi.nlm.nih.gov/29568823/ |
| **UNC-CH Neonatal Atlas (Saghafi et al. 2017)** | 30 (NN male/female) | DW | 2 (FA and average intensity maps) | 1 | 2×2×2 $mm^3$ | 14 days | Atlas was constructed with a patch-based method, that jointly considers neighbouring gradient directions in the DW images. A group regularization framework is used to constrain local patches for consistent spatio-angular reconstruction. |

| | | | | | | | |
|---|---|---|---|---|---|---|---|
| | | | | | | | https://pubmed.ncbi.nlm.nih.gov/28345171/ |
| **UNC Cortical (Li et al., 2015)**<br><br>https://bbm.web.unc.edu/tools/ | 35 individuals (18 males, 17 females); 202 scans (4–7 per infant; the number of scans was N = 35 at 1 month, N = 28 at 3 months, N = 31 at 6 months, N = 27 at 9 months, N = 29 at 12 months, N = 31 at 18 months, and N = 21 at 24 months | Structural: T1 and T2; DW | No anatomical parcellation provided | 7 | T1: 1×1×1 $mm^3$, 163 842 vertices across the cortical surfaces<br><br>T2: 1.25×1.25×1.95 $mm^3$ (resampled to 1×1×1 $mm^3$)<br><br>DW: 2×2×2 $mm^3$ (resampled to 1×1×1 $mm^3$) | 1, 3, 6, 9, 12, 18, and 24 months | Volumetric segmentation in line with their prior work (Li et al., 2013), i.e. a longitudinally consistent tissue segmentation by an infant-dedicated, 4D level-set method referencing iBEAT software (Wang et al., 2011, 2012, 2014). Groupwise surface registration was used for the creation of the cortical surface atlas with Spherical Demons (Yeo et al., 2010).<br><br>https://pubmed.ncbi.nlm.nih.gov/25980388/ |

| | | | | | | | |
|---|---|---|---|---|---|---|---|
| **UNC detail-preserved longitudinal 0-3-6-9-12 months-old atlas (Zhang et al., 2016)**<br><br>https://www.nitrc.org/projects/infant_atlas_4d/ | 35 (18 males, 17 females) | structural T1, T2 | 3 (GM, WM, CSF) | scanned every 4 months | T1: 1×1×1 $mm^3$<br><br>T2: 1.25×1.25×1.95 $mm^3$ (resampled to 1×1×1 $mm^3$) | Term-born, 0-1 years | Tissue segmentation was carried out with iBEAT. The template construction was carried out with a novel framework for consistent spatial-temporal construction of longitudinal atlases where the atlas construction was performed in spatial-temporal wavelet domain simultaneously.<br><br>https://pubmed.ncbi.nlm.nih.gov/27392345/ |
| UNC/UCI neonate hippocampus amygdala multi-atlas<br><br>https://www.nitrc.org/projects/unc_brain_atlas/ | 6 | Structural T1, T2 | 7 | 1 | T1; T2: 1×1×1 $mm^3$ | Term-born, 0-5 weeks | Manual segmentation protocol.<br><br>*Publication N/A* |
| UNC/UMN Baby Connectome Project (BCP) Atlases (Ahmad et al., 2023) | 37 (17 males, 20 females) | T1, T2 | | 25 (2 weeks + 1-24 months), infants were | T1; T2: 0.8×0.8×0.8 $mm^3$ | | Tissue segmentation using iBEAT V2.0 (https://ibeat.wildapricot.org). The 12-month surface-volume atlas was constructed using a Dynamic Elasticity Model with Surface Constraint (SC-DEM) for |

| | | | | | | | |
|---|---|---|---|---|---|---|---|
| | | | | divided into 6 cohorts, not all of were scanned at the same time points. | | | groupwise registration of tissue segmentation maps. The 2 weeks – 24 months longitudinal atlases were constructed using parallel transported longitudinal deformations.<br><br>https://pubmed.ncbi.nlm.nih.gov/36585454/ |
| **UNC volumetric/UNC-infant-0-1-2-atlases (Shi et al., 2011)**<br><br>http://bric.unc.edu/ideagroup/free-softwares/unc-infant-0-1-2-atlases/ | 95 same individuals at each time point (56 males, 39 females) | Structural: T2 for neonates and T1 for 1- and 2-year-olds | Labels drawn from Automated Anatomical Labeling (AAL) map (Tzourio-Mazoyer et al. 2002) with cortical and subcortical 90 labels | 3 | T1: $1\times1\times1$ $mm^3$<br><br>T2: $1.25\times1.25\times1.95$ $mm^3$ | 0, 1 and 2 years | ITK-SNAP (Yushkevich et al., 2006) was used for ground-truth manual segmentation of the neonates. SPM5 is used for atlas-based segmentation. A groupwise registration algorithm (Wu et al., 2011) was used for the atlas construction of each three age groups.<br><br>https://pubmed.ncbi.nlm.nih.gov/21533194/ |
| **USC atlas/template (Sanchez et al., 2012)** | Scan images obtained | NIHPD-Structural: T1 and T2 (only | 3 (GM, WM, CSF) | 13 | NIHPD-Structural: T1 and | Range 8 days - 4.3 years (13 | FSL FLIRT was used to make a preliminary template of four 6-month-olds' heads and brains from |

| | | | | | | | |
|---|---|---|---|---|---|---|---|
| http://jerlab.psych.sc.edu/neurodevelopmentalmridatabase | from two sources: NIHPD & MCBI; NIHPD = 105 (59 males, 46 females); MCBI = 49 (24 males, 25 females) | axial images acquired); MCBI-Structural T1 (sagittal plane) and T2 (axial plane) | | | T2 (only axial images acquired); MCBI-Structural T1 (sagittal plane) and T2 (axial plane) | groups; mean ages 2 weeks, 3, 4.5, 6, 7.5, 9, 12, 15, 18 months, 2, 2.5, 3, 4 years) | the USC-MCBI dataset; SPM8; and ANTS were used for template construction.<br><br>https://pubmed.ncbi.nlm.nih.gov/21688258/ |
| **Zhang DTI Atlas (Zhang et al. 2014)** | 9 (2 males, 7 females) | T1, DTI | 122 | 1 | T1: 1 mm slice thickness<br><br>DTI: 2 mm slice thickness | 2-13 days | The template was constructed using a volume-based template estimation (VTE) method. VTE was morphed to the JHU-neonate-SS atlas parcellation to label the anatomical features.<br><br>https://pubmed.ncbi.nlm.nih.gov/25026155/ |
